# The Unexpected Match: STAT3-NORAD Interaction, a Novel Link in Antiviral Defense

**DOI:** 10.1101/2023.06.29.546999

**Authors:** Amir Argoetti, Dor Shalev, Galia Polyak, Noa Shima, Hadas Biran, Tamar Lahav, Tamar Hashimshony, Yael Mandel-Gutfreund

## Abstract

In the intricate biological landscape, evolution often repurposes familiar elements for novel roles. This study reveals an interaction between the long non-coding RNA NORAD, noted for its role in DNA stability, and the immune-related transcription factor STAT3. Our findings indicate that NORAD’s binding to STAT3 facilitates its nuclear entry, suppressing the antiviral response. In the absence of NORAD, STAT3 remains cytoplasmic, enabling STAT1 to activate this response. Evidence from viral infections and clinical samples reinforces this unique role for NORAD. Intriguingly, while other functions of NORAD are conserved in evolution, this newly discovered role is unique to humans, owing to the introduction of an ALU element in hominoids. This discovery sheds new light on the evolution of antiviral defenses.

## Introduction

Transcription factors (TFs) are key regulators of gene expression that activate or repress transcription by binding to DNA. In recent years, it has become apparent that the activity of TFs can be mediated by RNA, specifically long non-coding RNAs (lncRNAs) (1–11). One such TF is Signal Transductor and Activator of Transcription 3 (STAT3), which was found to associate with lncRNAs in diverse human and murine cell types(8–11). STAT3 is known for its different roles in inflammation, the immune system, embryonic development, and cancer. In a recent study, we found that STAT3 interacts with a small, unique subset of RNAs in the cytoplasm of human embryonic stem cells (hESCs)(12). Among the identified targets was the Non-coding RNA Activated by DNA Damage (NORAD). This highly conserved, cytoplasmic lncRNA is most known for maintaining DNA stability. It does so by binding the RNA-binding protein (RBP) Pumilio at multiple sites across its transcript, through Pumilio Recognition Elements (PREs)(13–15). This sequesters Pumilio into phase-separated compartments termed RNA-protein (RNP) granules(16). In addition, an interaction between the 5’ region of NORAD and the RBP RBMX was shown to play a role in maintaining genome integrity(17). While the PREs and 5’ regions of NORAD have been extensively studied, not much is known about the function of other regions in the NORAD transcript, specifically the 3’ end, where we identified the interaction with STAT3(12).

## Results

### Knockdown of NORAD mimics viral response

Consistent with previous studies(13, 14) and our recent work suggesting that STAT3 interacts with NORAD outside of the nucleus, we found that NORAD localizes mostly to the cytoplasm of hESCs (fig. S1A). To test the possible function of its interaction with STAT3, we knocked down (KD) NORAD in hESCs. The downregulation was confirmed by RT-qPCR (fig. S1B(and its impact was assessed with RNA sequencing (RNA-seq). RNA-seq analysis revealed a significant upregulation of 29 genes (Padj<0.05, FC>2), 16 of which are Interferon Stimulated Genes (ISGs), including ISG15, IFI6, MX1, IFITs, and OASs. Gene ontology (GO) analysis reinforced that NORAD KD activates the “defense response to viruses” pathway (Fig. 1A and Table S1). To explore whether the response is unique to pluripotent cells, we repeated the NORAD KD and following assays in fully differentiated foreskin fibroblasts (HFFs) (fig. S1B(. In this case, it resulted in an even more robust upregulation of the antiviral response, observed from eight hours post-transfection and most prominently at 24 hours (Fig. 1B, C and Table S2).

**Figure 1:**
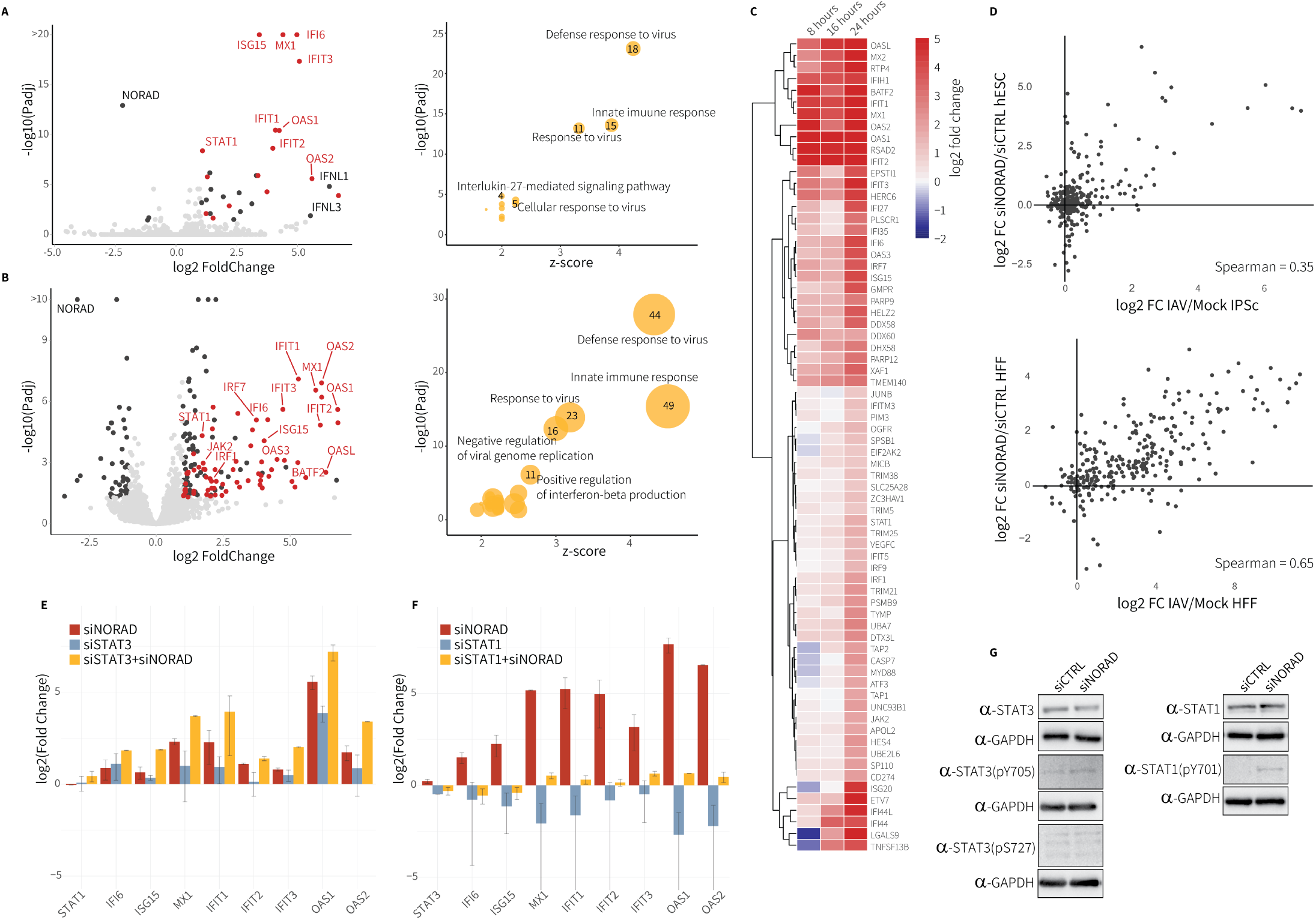
Knockdown of NORAD in pluripotent and differentiated human cells actives the innate immune response via the STAT1 regulatory pathway. **(A and B)** Left: Volcano plot of RNA-seq data from NORAD KD versus control siRNA. Differentially expressed (Padj<0.05, -2>FC>2) ISGs are marked in red. Right: Gene ontology enrichment analyses of the DEGs upon NORAD KD. **(A)** hESCs 72 hours post siRNA transfection (n=4) **(B)** HFFs 24 hours post transfection (n=4). **(C)** heatmap of FCs of ISGs 8, 16, 24 hours post transfection in HFFs, shown are only differentially expressed ISGs in at least one time point **(D)** Scatterplots demonstrating the correlations between FCs calculated for all expressed ISGs in IAV relative to mock-infected cells and the FCs calculated for ISG expression in NORAD KD in this study. Up: Comparison between IAV infection in hIPSCs to siNORAD treatment in hESCs. Down: Comparison between IAV infection to NORAD KD in HFFs. **(E-F)** RT-qPCR analysis of ISGs expression in double KD experiments in HFFs of STAT3+NORAD **(E)** and STAT1+NORAD **(F)**. siNORAD (red), siSTAT (light blue) and siSTAT + siNORAD (yellow) relative to siCTRL. **(G)** Western blot analysis of STAT3 and STAT1 protein expression in HFFs 24 hours post siRNA transfection.

The number of ISGs that were induced in response to the KD in HFFs was significantly higher than in hESCs (69 genes, Padj<0.05, FC>2) (Fig. 1B). When analyzing RNA-Seq data from HCT116 cells in which NORAD was downregulated by CRISPRi(17), a similar upregulation of ISGs is observed at 24 hours post-induction. Of note, the interferon genes themselves were not upregulated in any of these conditions (Fig. 1B) (17). Remarkably, the changes in ISGs expression in the knocked-down cells correlate with the changes in the same cell types infected by influenza A virus (IAV), though this correlation was weaker in hESCs (Fig. 1D) (data from Egenberger et al.(18)). Moreover, the activation of the antiviral response in both IAV-infected and NORAD KD in HFFs was noticeably higher than in hESCs (Fig. 1D). This can be explained by the fact that ESCs do not utilize the canonical interferon immune response – rather, they have an alternate pathway based on RNAi (18). In addition, consistent with previous findings(19), we found differences in the basal expression levels of a subset of ISGs between the non-treated HFFs and hESCs (fig. S1C). Nevertheless, in the absence of NORAD, ISGs were activated in both pluripotent and differentiated cells (Fig. 1A, B). TF enrichment analysis using ChEA3(20) strongly evoked the TFs STAT1-3 as the regulators of the observed gene expression changes upon NORAD KD (fig. S1D, E). This suggests a functional role to the STAT3-NORAD interaction in the transcriptional circuit regulating the interferon pathway.

### NORAD affects STAT1/STAT3 balance in the nucleus

Next, we tested the effect of NORAD KD on STAT3 and STAT1 protein level, modifications, and localization. Overall, the levels of the transcriptionally active form of STAT3 (p-Y705) slightly increased, while the total STAT3 protein level and its transcriptionally enhanced form (p-S727) did not considerably change (Fig. 1G). On the other hand, overall STAT1 protein levels and specifically STAT1 (p-Y701), a modification associated with nuclear translocation and transcription activation(21), increased substantially (Fig. 1G). When looking at their cellular localization, we observed a significant shift upon NORAD KD – STAT3 was less present in the nucleus, whereas STAT1 nuclear levels increased (Fig. 2, A-C and fig. S2). Unlike STAT1, STAT3’s nuclear localization could not be explained by changes in its phosphorylation. An alternative path requires the interaction of unphosphorylated STAT3 with the Importin-β1 (KPNB1) (22). In a co-immunoprecipitation assay, we found that NORAD mediates the interaction between the two (Fig. 2D). Our results are also supported by data from recent studies, where Importin-β1 was identified in the interactomes of NORAD that used its 3’ region as bait (13, 14).

**Figure 2:**
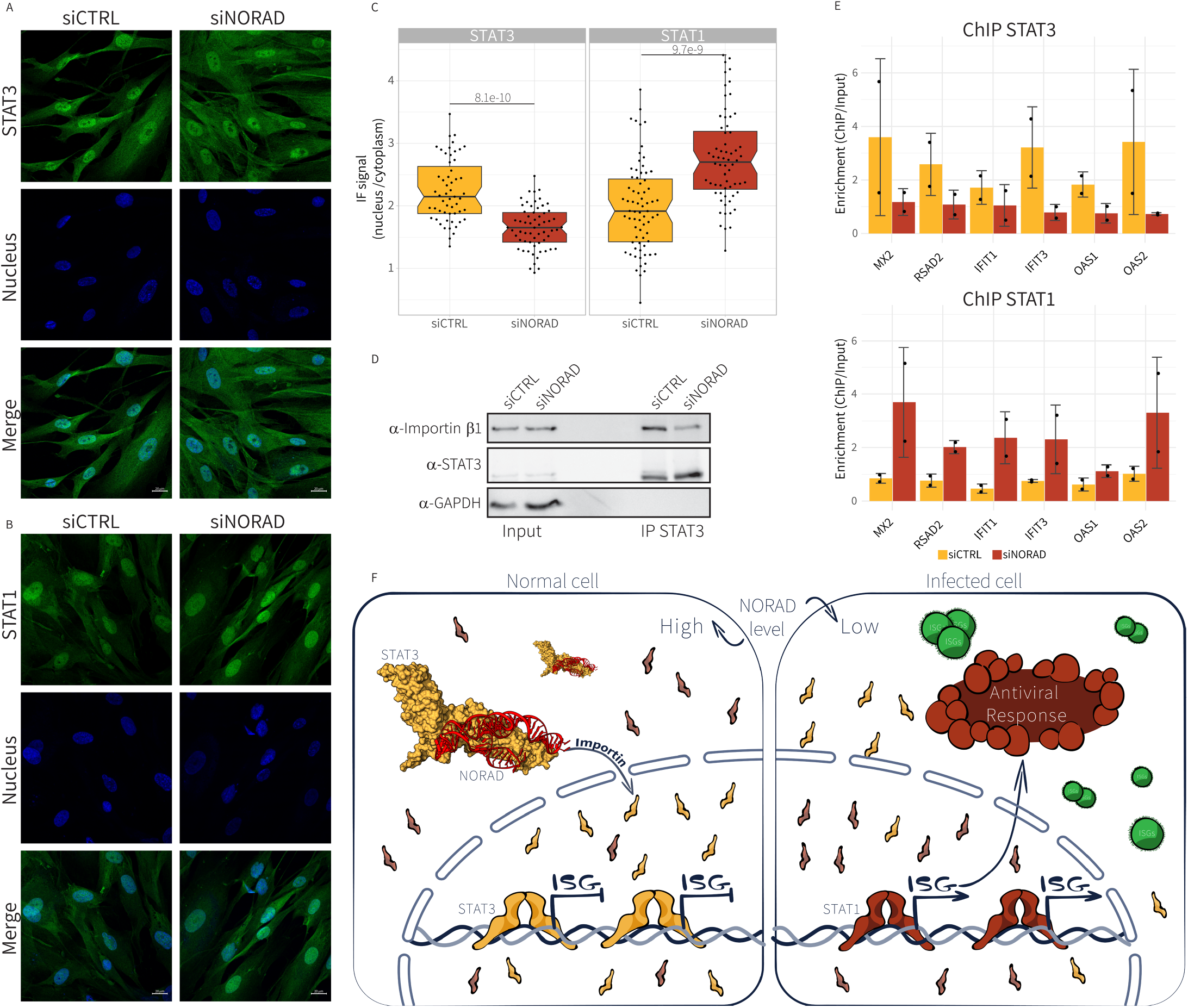
The activation of the innate response upon NORAD KD is mediated by STAT1-STAT3 pathway. **(A and B)** An immunofluorescence visualization - upon NORAD knockdown compared to siCTRL samples in HFFs (16h post siRNA transfection), **(A)** STAT3 **(B)** STAT1 (In blue: nuclear staining by DAPI). **(C)** Box plots of the ratio between the nucleus and cytoplasm IF signal (55 and 61 cells for STAT3 siCTRL and siNORAD, respectively and 71 and 64 cells for STAT1 siCTRL and siNORAD, respectively) showing significant differences between the different conditions (STAT3 p-value = 8.1 e-10, STAT1 p-value = 9.7e-9, Mann Whitney Wilcoxon test). **(D)** Western blot analysis of Co-IP experiment. HFF lysates from siNORAD and siCTRL samples immunoprecipitated by anti-STAT3. Protein-protein interactions were immunodetected by anti-Importin-β1, anti-STAT3 and anti-GAPDH. **(E)** Binding enrichment of STAT3 (top) and STAT1 (bottom) to the promoters of six selected ISGs in siCTRL and siNORAD in HFFs. DNA binding levels were determined by ChIP-qPCR (n=2). **(F)** Illustration of the proposed model for the role of NORAD-STAT3-interaction in regulating ISG transcription.

We next sought to examine whether the change in the ratio between STAT3 and STAT1 in the nucleus influences their binding preference. We carried out Chromatin Immunoprecipitation (ChIP)-qPCR in HFFs, both with and without NORAD KD. We focused on the promoter regions of a panel of ISGs (MX2, RSAD2, IFIT1, IFIT3, OAS1, OAS2), which are known targets of STAT1 and STAT3. Following NORAD KD, our ChIP results revealed a significant decrease in the occupancy of STAT3 at the tested ISGs. Conversely, the binding of STAT1 to the same promoters was significantly increased in the KD cells (Fig. 2E). This observed change in the binding of STAT1 and STAT3 in response to NORAD KD is consistent with the upregulation of these ISGs. Altogether, we show that in the presence of NORAD, STAT3 interacts with Importin-β1 and is translocated to the nucleus, where it binds to ISG promoters and inhibits their transcription. Upon NORAD down-regulation, STAT3’s nuclear levels decrease, allowing STAT1 to bind in its place and activate the interferon pathway. Early activation of these ISGs increases the phosphorylation levels of STAT1, thereby enhancing the transcription of both these initially activated ISGs and many others. Together, this results in positive feedback that intensifies the antiviral response. (Fig. 2F)

### NORAD is downregulated in viral infections

Building on our finding that NORAD downregulation triggers the antiviral response, we aimed to gauge NORAD’s involvement in viral infections. To begin, we conducted a general assessment of lncRNA expression in cells infected by three different viruses: SINV(23), SARS-CoV-2(24) (RNA viruses), and HSV1(25) (DNA virus). We found 928 lncRNAs whose expression changed in at least one dataset, 60 of which changed in at least two, and only six were differentially expressed in all three infections. Out of these six lncRNAs, four were upregulated: SNHG1, SNHG12, SNHG15, and LINC00511, and two were downregulated: NORAD and its pseudogene HCG11 (14) (Fig. 3A, Table S3). We next examined the effect of viral infection on NORAD based on clinical data. We analyzed single-cell RNA-seq data from nasopharyngeal samples of four COVID-19 patients compared to three healthy controls(26), and found a significant decrease of NORAD in epithelial cells (Fig. 3B, C). A consistent significant reduction was also observed by Butler et al. (27) in bulk RNA-seq data from a larger cohort of 173 COVID-19 positive patients relative to the expression in 407 negative controls (nasopharyngeal samples).

**Figure 3:**
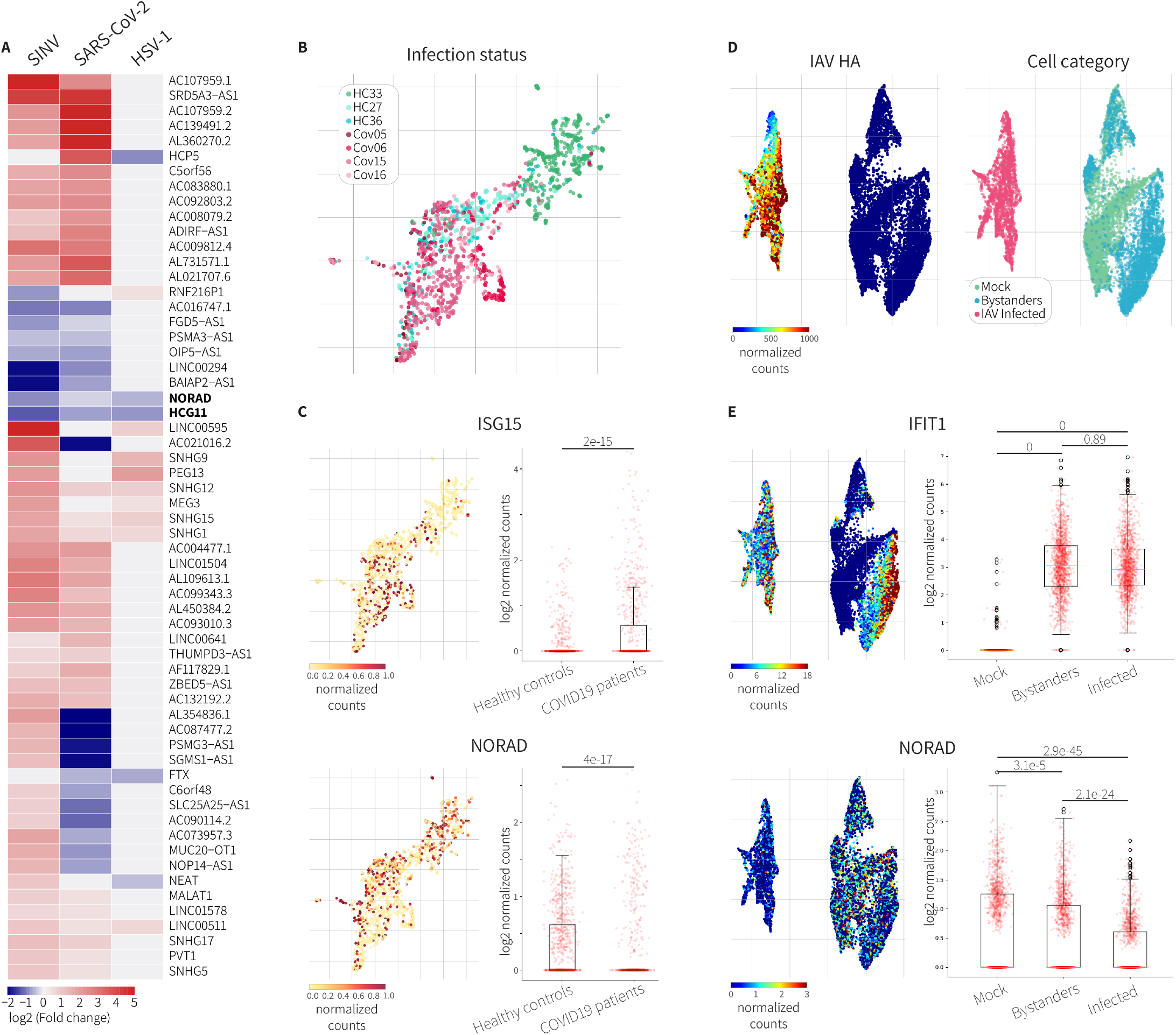
NORAD lncRNA is down regulated in viral infections. **(A)** Heatmap of differentially expressed lncRNAs upon infection in SINV, SARS-CoV-2 and HSV1. **(B-C)** Analyses of NORAD expression levels in COVID-19 patients compared to healthy controls **(B)** UMAP colored by the patient label (green and red shades for Healthy control (HC) and COVID patients (Cov), respectively). **(C)** UMAP colored by ISG15 (top) and NORAD (bottom) levels. Quantification of the gene expression levels are demonstrated in boxplots besides each UMAP. P-adj values are depicted above the plots (Mann Whitney Wilcoxon test). **(D-E)** Analysis of NORAD expression in cells infected by IAV compared to mock infected cells, based on single cell RNA-seq data. **(D)** Left: UMAP colored by hemagglutinin viral gene IAV-HA. Right: UMAP colored by the cell category. **(E)** UMAPs colored by IFIT1 (top) and NORAD (bottom) levels. Quantification of the expression levels of the two factors are demonstrated in boxplots besides each UMAP. P-adj values are depicted above the plots (Mann Whitney Wilcoxon test). As shown NORAD levels are significantly decreased in the virus infected cells compared to mock and bystanders.

The downregulation of NORAD upon viral infection could be explained in two ways. Either it is caused directly by the penetration of the virus into the cell, or NORAD could be downregulated as part of the interferon pathway response. To address this question, we analyzed single-cell RNA-seq data from human lung cells infected by Influenza-A virus or a mock infection (28). Based on the experimental setup, we classified the cells into two categories: ‘mock’ for control cells and ‘exposed’ for those the virus Introduced to the culture. The exposed cells were then subdivided into two categories based on viral gene expression, as defined in Ramos et al.(28): ‘infected’ cells and ‘bystander’ cells - uninfected but responsive to interferon secretion by the adjacent infected cells (Fig. 3D).

While both infected and bystander cells express similar high levels of ISGs, NORAD was significantly reduced in the infected cells compared to mock and bystanders (Fig. 3E and fig. S3). These observations suggest that the downregulation of NORAD is likely a direct cellular response to the presence of the virus, rather than to the Interferon.

### A novel function for NORAD in evolution

The role of NORAD lncRNA in DNA stability has been revealed to be highly conserved in mammals (13–15). Several lncRNAs are known to play a role in regulating the immune response, both in humans and mice(29–32). Upon demonstrating the involvement of NORAD in the defense against viruses, we were intrigued as to how this novel function is conserved in mammals. To address this, we knocked down Norad in mouse embryonic fibroblasts (MEFs), followed by RNA-seq (fig. S4A, B and Table S4). While NORAD KD in human cells resulted in upregulation of ISGs (Fig. 4A), Norad downregulation did not activate the transcription of ISGs in murine cells (Fig. 4B and fig. S4A, B). Consistently, no interaction between Stat3 and Norad in MEF cells was detected in CLIP-qPCR (Fig. 4C and Fig. S4C). Evolutionary conservation analysis of NORAD lncRNA found a lower conservation of the STAT3 binding site identified in eCLIP(12) compared to the PREs associated with the NORAD repeats(14) (Fig. 4D). Interestingly, the STAT3 binding site is located at approximately 100 nts downstream to a primate-specific ALU element. Phylogenetic analysis of the region in which STAT3 binds NORAD in humans was performed on 30 mammals. This demonstrates that sequences containing an ALU element upstream to the STAT3 binding region fold to a similar stem-loop structure (Fig. 4E). This prediction is highly consistent with the experimentally solved structure of this region in human NORAD by COMRADES(33). LocARNA(34) multiple alignment of the binding region of STAT3 on NORAD from diverse species (extracted from 10 ALU-containing primates, five non-ALU-containing primates, three rodents, and 12 other mammals) propose that the observed differences in the predicted local structure between human and mouse may be related to the presence of the transposable element (fig. S4D). We propose that the insertion of an ALU in the 3’ region of NORAD during primate evolution induced a change in the structure of this region, enabling an interaction with STAT3. The novel interaction between STAT3 TF and NORAD lncRNA resulted in the birth of a new mechanism in evolution, which enables a rapid activation of the interferon pathway.

**Figure 4:**
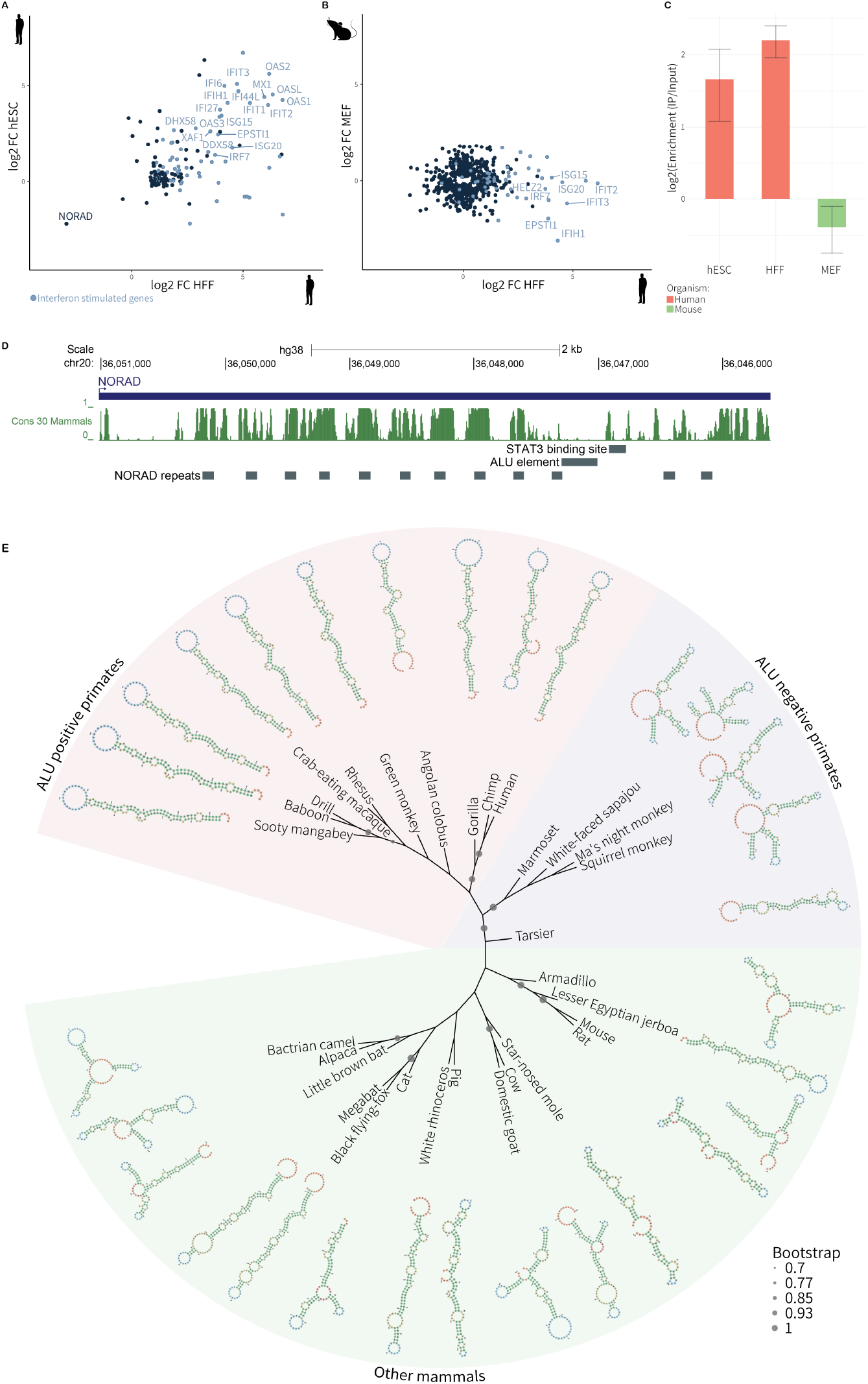
NORAD downregulation does not induce ISG expression in mouse cells consistent with the lack of conservation of the STAT3 binding site on its 3’ region. **(A and B)** The FCs of differentially expressed genes upon NORAD KD in HFFs compared to hESCs **(A)** and HFFs compared to MEFs **(B)** showing a high correlation between the undifferentiated and the fully differentiated human cells while no significant changes in gene expression in MEFs. **(C)** CLIP-qPCR validation of STAT3 binding to NORAD in hESCs, HFFs and MEFs. As demonstrated, there is no interaction between STAT3 and Norad in the mouse cells. **(D)** Evolutionary conservation across the NORAD transcript, depicting weak sequence conservation at the STAT3 binding site. STAT3 binding site, ALU and NORAD repeats are illustrated in grey rectangles **(E)** Phylogenetic tree based on the sequence of the STAT3 binding site (135 nts). RNA predicted structures (predicted by RNAfold) are presented. Bootstrap values above 0.7 (calculated based on 1000 trees) are represented in circles.

## Discussion

In this study, we have discovered a novel role for the lncRNA NORAD in regulating the innate immune response in human cells. These results are consistent with previous studies that have shown that other lncRNAs can act as regulators of the antiviral response through diverse mechanisms (29–32). The involvement of a lncRNA in sensing and transducing a signal enables a highly efficient and quick cellular response to viral infections, without the need for protein metabolism. Here we have found that NORAD is required for the translocation of the unphosphorylated STAT3 to the nucleus via Importin-β1, where it binds to the promoters of interferon-stimulated genes and negatively regulates their transcription. We also observed that NORAD levels are reduced in human viral infections, both in cells infected in-vitro and in patients. This can be explained by recent studies that show that among other RNAs, NORAD is recruited to stress granules upon viral infection, where it is cleaved by RNaseL (35–37) as part of the intrinsic cellular antiviral response. This is an extremely rapid reaction that inhibits viral replication without requiring changes in gene expression.

Our findings demonstrate that NORAD downregulation releases the STAT3-mediated inhibition of ISGs, thus activating the innate immune response by STAT1. Collectively, these results suggest that the lncRNA NORAD may serve as a bridge between the intrinsic defense response and the more established STAT1-mediated innate immune response. The link that NORAD generates between the two pathways may lead to a more direct transcriptional activation of ISGs, bypassing many regulatory processes. This is particularly significant given that many viruses have evolved strategies to inhibit the classical activation of the interferon pathway, a key element of the host’s antiviral defense. This ‘shortcut’, or non-canonical activation, could circumvent these viral inhibitions, as it does not rely on the activation of IRFs and interferon secretion. Such a mechanism might expedite the response of the host defense system, providing a rapid reaction to viral threats while still invoking the robustness of the STAT1-mediated innate immune response. It is interesting to note that pluripotent cells do not normally activate the interferon pathway. Therefore, the upregulation of ISGs induced by NORAD KD reinforces that the new role suggested for NORAD does not rely on interferon stimulation.

Unlike protein-coding genes, the majority of lncRNAs have evolved under little to no selective constraints(38). Thus, lncRNAs have the potential to better adapt to rapid changes in evolution compared to proteins. We found that the interaction between STAT3 and NORAD is mediated by a structured region in the 3’ domain of NORAD, downstream to the ALU element that is present only in hominoids. Based on a comparative RNA structure analysis of this region, we suggest that the introduction of an ALU element to NORAD in primates led to a structural change that was conserved in hominoids evolution. We propose that this enabled the binding of STAT3, giving rise to a new function for NORAD in regulating the antiviral response. It has been suggested that mammalian embryonic stem cells keep the interferon response silent as it interferes with their two defining characteristics; namely self-renewal and pluripotency (18, 19). Our identification of NORAD as the link between the interferon pathway and the intrinsic cellular response could provide a rationale for a reduced intensity of this intrinsic response in hESCs compared to mESCs. This may suggest a protective mechanism to mitigate the activation of the interferon response (39).

Together, we show that a non-classical interaction between a lncRNA and a bona fide TF plays a key role in the rapid cellular response to external stimuli, adding an additional important layer of regulation in eukaryotes.

## Methods

### Cell culture

Human foreskin fibroblasts (HFFs) and Mouse embryonic fibroblasts (MEF) were cultured in Dulbecco’s modified eagle medium (DMEM) supplemented with 10% fetal bovine serum (FBS), glutamine, and penicillin/streptomycin (pen/strep). The cells were maintained in a humidified incubator at 37°C with 5% CO2 and passaged every 3-4 days by trypsinization and re-seeding at a split ratio of 1:3. Human embryonic stem cells (hESCs) were maintained in suspension in YF100 medium as described in (12). The hESCs were split into single cells by treatment with EGTA and TrypLE and then transferred to Matrigel-coated plates in mTeSR1 medium supplemented with 5 μM ROCK inhibitor. The cells were allowed to attach to the plates for at least 24 hours before the start of the experiment.

### Single-molecule FISH

Cells were fixed using 4% formaldehyde in PBS for 15 minutes at room temperature, followed by a rinse with 1X PBS. After rinsing, the samples were incubated with cold 70% ethanol at 4°C for at least 2 hours. Subsequently, cells were washed twice with wash buffer (10% Formamide, 2X SSC) for 5 minutes each. The probes, labeled with Quasar® 670 Dye, were diluted in UPW to a working solution of 2.5μM. A volume of 5μl of these probes was added to 45μl of hybridization buffer (10% Dextran sulfate, 10% Formamide, 1mg/ml E.coli tRNA, 2X SSC, 0.02% BSA, 2mM Vanadyl-ribonucleoside complex). This mix was applied onto a clean piece of parafilm for each sample.

The samples were carefully placed section-down onto a drop of the hybridization mix and incubated overnight at 37°C. The next day, samples were washed once in wash buffer for 30 minutes at 37°C. Concurrently, a solution of wash buffer with 1:200 of Dapi (10μg/ml) was pre-warmed for 30 minutes together with the samples.

The samples were then incubated in GLOX buffer, allowed to stand for a few minutes, mounted with Prolong Gold Antifade Mountant (life technologies, CAT: P36930), and immediately visualized on a Leica DMI8 inverted fluorescent microscope with an x68 oil-immersion objective. The dot detection was carried out automatically using Imaris software.

### NORAD knockdown

HFF and hESC and MEF were transfected with small interfering RNAs (siRNAs) using Lipofectamine RNAiMAX (Invitrogen). TriFECTa kit DsiRNA Duplex (IDT) containing three RNAi duplexes were designed to specifically target the NORAD lncRNA. The cells were incubated with the siRNAs for 8,16, and 24 hours for HFF, 24 hours for MEF, and 72 hours for hESC. After the incubation period, total RNA was purified using TRI-reagent (Sigma).

To evaluate the knockdown efficiency, qRT-PCR was performed using primers specific for the NORAD lncRNA and the reference gene GAPDH.

STAT1 and STAT3 knockdown was performed in HFFs by RNAi as described above. After transfection cells were incubated for 72 hours. For double knockdown, cells were transfected with siSTAT1/3 and after 48 hours were further transfected with siNORAD.

### RNA sequencing

RNA sequencing was performed using the CEL-seq2 protocol(40) and sequenced on an Illumina HiSeq 2500 platform.

To analyze the CEL-seq2 RNA sequencing data, the reads were first processed using the cutadapt software (41), then aligned to the reference genome using RNA-STAR (42). HTSEQ count (43) was used to generate raw count data, and the DESeq2 R package (44) was used to conduct differential gene expression analyses. GO enrichment was analyzed using DAVID web server (https://david.ncifcrf.gov/summary.jsp) and visualized by the GOplot R package (45)

### ChIP-qPCR

Chromatin immunoprecipitation (ChIP) followed by quantitative PCR (qPCR) was performed to examine the DNA binding of STAT1 and STAT3. Cells were crosslinked with 1% formaldehyde for 10 minutes at room temperature and quenched with 125 mM glycine. Chromatin was lysed in FA lysis buffer (50mM HEPES-KOH pH7.5, 140mM NaCl, 1mM EDTA, 1% Triton X-100, 0.1% sodium deoxycholate, 0.1% SDS, 1mM PMSF) and sonicated by Sonics Vibracell VCX130 for 2.5 minutes (10 pulses of 15’’ on 10’’ off). Immunoprecipitation was performed using 3μg antibody against STAT1 or STAT3 (Cell Signaling Cats:#14994, #12640 respectively) for 4 hours and afterward incubated with 50 μl Protein G magnetic beads (Invitrogen Cat:#10004D). Beads were washed in RIPA buffer (50Mm Tris-HCl pH8, 150mM NaCl, 2Mm EDTA, 1% NP-40, 0.5% sodium deoxycholate, 0.1 SDS, 1mM DTT, 1mM PMSF) and the immunoprecipitated DNA was eluted with proteinase K and purified by MN NucleoSpin Gel and PCR cleanup kit. qPCR was performed using primers specific to the ISGs and ACTB The qPCR reactions were performed in triplicate, and the enrichment of ISGs or ACTB was calculated as the percent input after normalization to the total input DNA.

### Western blot analysis

Total protein was extracted from cells using iCLIP lysis buffer (50 mM Tris-HCl pH 7.5, 100 mM NaCl, 1% NP-40, 0.5% sodium deoxycholate, 0.1% SDS, 1mM PMSF, 1% Phosphatase inhibitors cocktail 2 Sigma-Aldrich Cat:P5726, 1% Phosphatase inhibitors cocktail 3 Sigma-Aldrich Cat:P0044). Western blot analysis was performed using antibodies against STAT1, STAT3, phosphorylated STAT1 (pSTAT1) at residue 701, and phosphorylated STAT3 (pSTAT3) at residues 705 and 727 (Cell Signaling Cats:#14994, #12640, #7649,#9145,#9134 respectively). GAPDH (Abcam Cat: ab8245) was used as a loading control.

### Immunofluorescence

Immunofluorescence staining was performed to visualize STAT3 and STAT1 proteins in hESCs and HFF cells. Cells were fixed in 4% formaldehyde for 15 minutes at room temperature. Cells were blocked and permeabilized with 5% normal goat serum and 0.3% Triton X-100 in PBS for 1 hour at room temperature and incubated with primary antibodies against STAT1 or STAT3 (Cell Signaling Cats:#14994, #12640 respectively) overnight at 4°C. Cells were washed with PBS and incubated with alexafluor 488-conjugated secondary antibody (Abcam Cat: ab150077) for 1 hour at room temperature. Nuclei were stained with DAPI for 5 minutes at room temperature. Cells were mounted with Fluoromount-G (Thermo Fisher Scientific Cat: 00-4958-02) and visualized using an LSM700 microscope (Zeiss). Images were analyzed using Fiji software. The nuclear to cytoplasm ratio was calculated for each cell individually by measuring the average signal inside ROIs.

### Co-Immunoprecipitation

Co-IP was performed to analyze the interaction between STAT3 and Importin-β1 in HFFs. Cells were crosslinked with 1% formaldehyde for 10 minutes at room temperature and quenched with 125 mM glycine. Cells lysed in FA lysis buffer (50mM HEPES-KOH pH7.5, 140mM NaCl, 1mM EDTA, 1% Triton X-100, 0.1% sodium deoxycholate, 0.1% SDS, 1mM PMSF) then Immunoprecipitation was performed using an antibody against STAT3 (Cell Signaling Cat:#12640) and Protein G magnetic beads (Invitrogen Cat:#10004D). The immunoprecipitated proteins were eluted from the beads with NuPAGE LDS sample buffer (Invitrogen Cat:NP0007) and analyzed by western blotting using an antibody against Importin-β1 (abcam Cat: ab2811). GAPDH (Abcam Cat: ab8245) was used as a loading control.

### CLIP-qPCR

Cells from hESCs, HFF, and MEF underwent UV cross-linking at 254 nm (3.2J/cm2 for hESCs and 1J/cm2 for HFF and MEF) to stabilize RNA-protein interactions. Post-crosslinking, cells were lysed and DNase-treated for DNA digestion. The cell lysate was then subjected to immunoprecipitation using a STAT3 antibody (3μg) to isolate STAT3 and its associated RNAs. Post-immunoprecipitation, samples were washed and digested with Proteinase K to liberate bound RNA, followed by RNA purification by phenol-chloroform extraction and the Zymo RNA Clean & Concentrator kit.

The extracted RNA underwent reverse transcription using SuperScript III Reverse Transcriptase with random hexamers at 25°C for 5 minutes, then at 55°C for 50 minutes to produce cDNA.

qPCR analysis of the cDNA, utilizing NORAD-specific primers, was conducted with GAPDH serving as an internal control. The ΔΔCt method was used for data analysis, determining the enrichment of NORAD RNA in STAT3 immunoprecipitates relative to input samples.

### RNA-seq Data Analysis

Publicly available RNA-seq datasets from cells infected with three different viruses - SINV(23), SARS-CoV-2(24), and HSV1(25) - were reprocessed and analyzed. For each dataset, the raw counts were imported and handled using the DESeq2 R package(44) for both normalization and differential expression analysis at each timepoint relative to control samples. Differentially expressed genes were defined as those with an adjusted p-value (Padj) <0.05. We chose the most responding timepoint as the one with the highest number of significantly upregulated ISGs. The differentially expressed lncRNAs identified in each dataset were then compiled together to identify lncRNAs that were differentially expressed on two out of three datasets.

### Single-cell RNA-seq Data Analysis for COVID-19 Patients

Single-cell RNA-seq data were analyzed from Seq-Well libraries of nasal wash samples from COVID-19 patients, generated by Gao et al.,(26). Raw count data was filtered to eliminate potential dead cells and doublets by removing cells with an atypical number of detected genes (Specifically-less than 500 or more than 10,000) (46). We also removed cells in which more than 33% of the reads were mapped to mitochondrial genes, since this may indicate cell stress (47, 48). Counts from the remaining cells were then normalized to the median counts of all cells.

Epithelial cells were identified based on gene expression criteria (KRT7>=1 and VIM<3). Within these epithelial cells, the expression levels of NORAD and ISG15 were compared between COVID-19 patients and healthy controls using a Mann-Whitney U test.

### Single-cell RNA-seq Analysis of Influenza A Virus-infected Cells

For the analysis of 10x Genomics single-cell RNA-seq data from Influenza-A virus and mock-infected human lung cells (28) we adapted the Python pipeline previously used for COVID-19 patient samples with a few modifications.

We first filtered cells that had less than 2500 or more than 5000 identified genes. Then, normalization was performed as described above, using median counts across all cells.

The cells were then classified into three groups:

1) Mock-infected cells defined as in the original sequencing library.

2) Infected cells expressing the Influenza A virus PR8_NP gene above 90 reads

3) Bystander cells that do not express the PR8_NP (below 90 reads).

Differential expression analysis was performed using a Mann-Whitney U test, and p-values were corrected for multiple testing using Bonferroni correction.

### Phylogenetic Analysis and Secondary Structure Prediction

Sequences corresponding to the STAT3 binding site in human (hg38 chr20:36,046,781-36,046,915) were extracted using the UCSC Genome Browser. For primates, sequences were derived from the Mammals Multiz Alignment & Conservation (27 primates), for other mammals, sequences were obtained from the Vertebrate Multiz Alignment & Conservation (100 Species).

Sequences extracted from the reverse strands were aligned using the T-Coffee web server (https://www.ebi.ac.uk/Tools/msa/tcoffee/). A phylogenetic tree was subsequently built using the BioNJ algorithm(49), with confidence assessed through 1000 bootstrap replicates. The resulting tree was visualized using the iTOL tool(50).

Simultaneously, the RNA secondary structure of the STAT3 binding site was predicted using the Minimum Free Energy (MFE) function in RNAfold (http://rna.tbi.univie.ac.at/cgi-bin/RNAWebSuite/RNAfold.cgi). The predicted structures were visualized in Forna (http://rna.tbi.univie.ac.at/forna).

The final assembly of the figure, which included both the phylogenetic tree and predicted RNA structures, was conducted in Adobe Illustrator.

## Supporting information

Primers and oligos

## Acknowledgments

This work was supported by the Israel Science Foundation (ISF) grant # 1182/16 and grant # 1556/22. We thank Dr. Nitsan Fourier from the Technion Genome Center for help and advice with RNA-Seq experiments, Dr. Nitsan Dahan, and Dr. Yael Lupu-Haber from the Life Sciences & Engineering infrastructure center for support and aid with microscopy imaging. We thank Anna Argoetti for her help in generating the illustration in Figure 2F.

---

**Figure S1.**
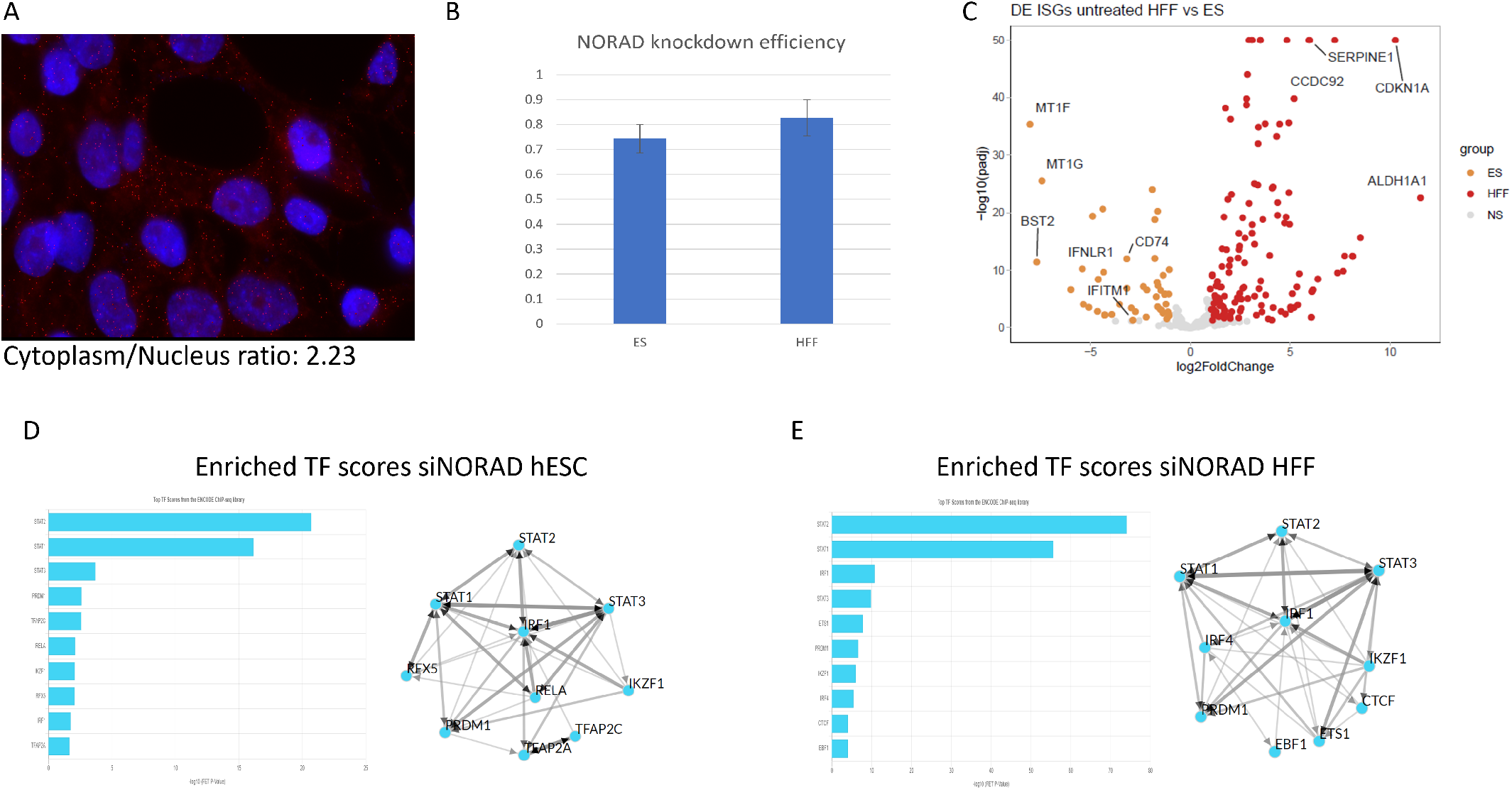
**(A)** Single-molecule Fluorescence In Situ Hybridization (smFISH) of the long non-coding RNA (lncRNA) NORAD in hESCs. The FISH data demonstrates that NORAD is predominantly located in the cytoplasm with a 2.2-fold higher concentration than in the nucleus. This suggests a significant role for NORAD in cytoplasmic processes. **(B)** NORAD knockdown efficiency in hESCs and HFFs, detected by rt-qPCR. **(C)** Volcano plot illustrating differentially expressed ISGs between hESCs and HFFs. The plot reveals a greater expression of ISGs in HFFs as compared to hESCs, indicating a differential baseline of the innate immune responses between these two cell types. **(D, E)** ChIP Enrichment Analysis (ChEA3) of differentially expressed genes upon knockdown of NORAD in (D) hESCs and (E) HFFs. Bar plots and network visualizations are provided. As shown, in both hESC and HFFs STAT1, STAT2, and STAT3 are at the top of the list of transcription factors that are predicted to regulate these differentially expressed genes. Results highlight the potential broad influence of NORAD on STAT-mediated signaling pathways.

**Figure S2.**
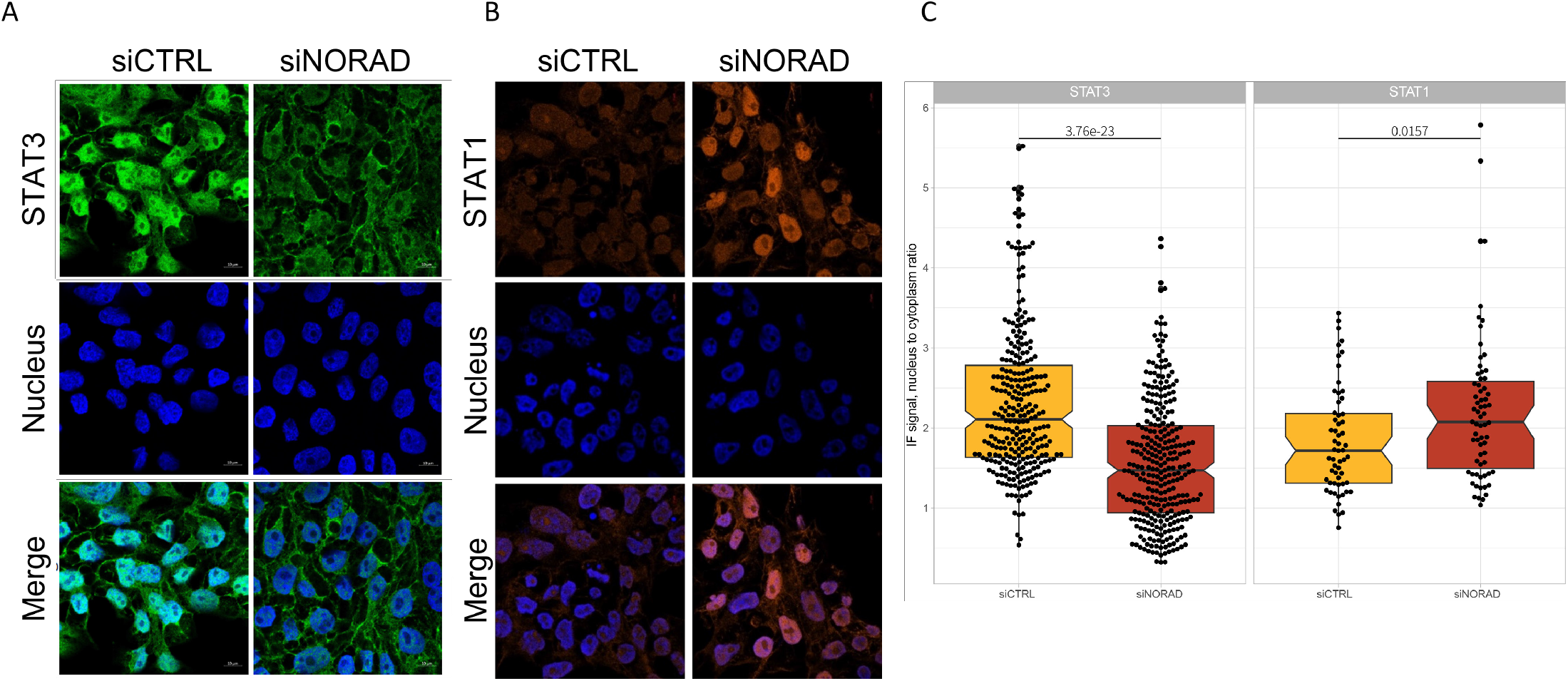
**(A**,**B)** An immunofluorescence visualization - upon NORAD knockdown compared to siCTRL samples in hESCs (72h post siRNA transfection), **(A)** STAT3 **(B)** STAT1 (In blue: nuclear staining by DAPI). **(C)** Box plots of the ratio between the nucleus and cytoplasm IF signal (282 and 302 cells for STAT3 siCTRL and siNORAD, respectively and 59 and 67 cells for STAT1 siCTRL and siNORAD, respectively) showing significant differences between the different conditions (STAT3 p<3.76e-23, STAT1 p<0.0157, Mann Whitney Wilcoxon test)

**Figure S3.**
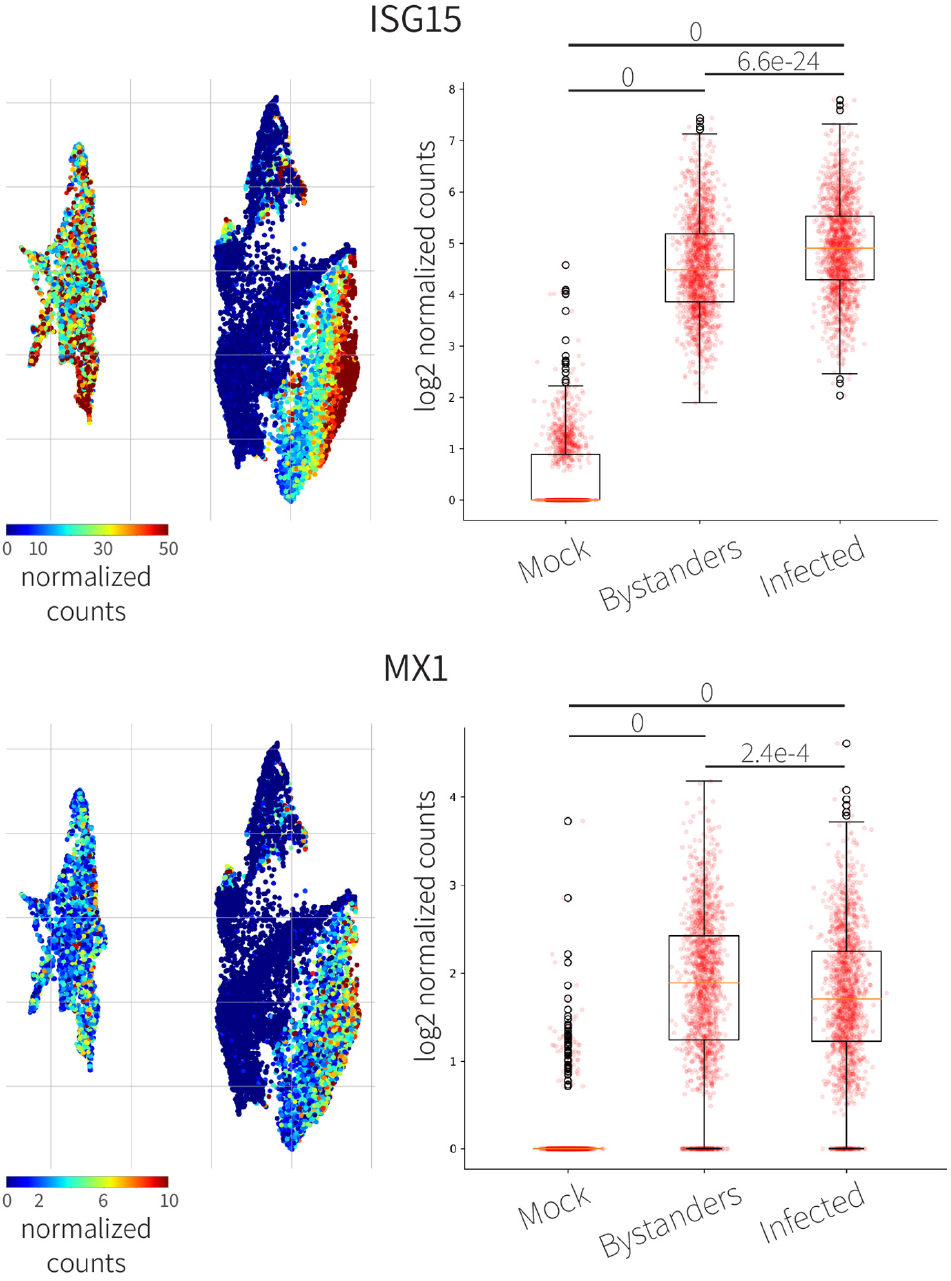
Single cell RNA seq analysis of IAV infected cells compared to mock infected cells, based on single cell RNA-seq data. UMAPs colored by ISG15 (top) and MX1 (bottom) levels. Quantification of the expression levels of the different factors are demonstrated in boxplots besides each UMAP P-adj values are depicted above the plots (Mann Whitney Wilcoxon test)

**Figure S4.**
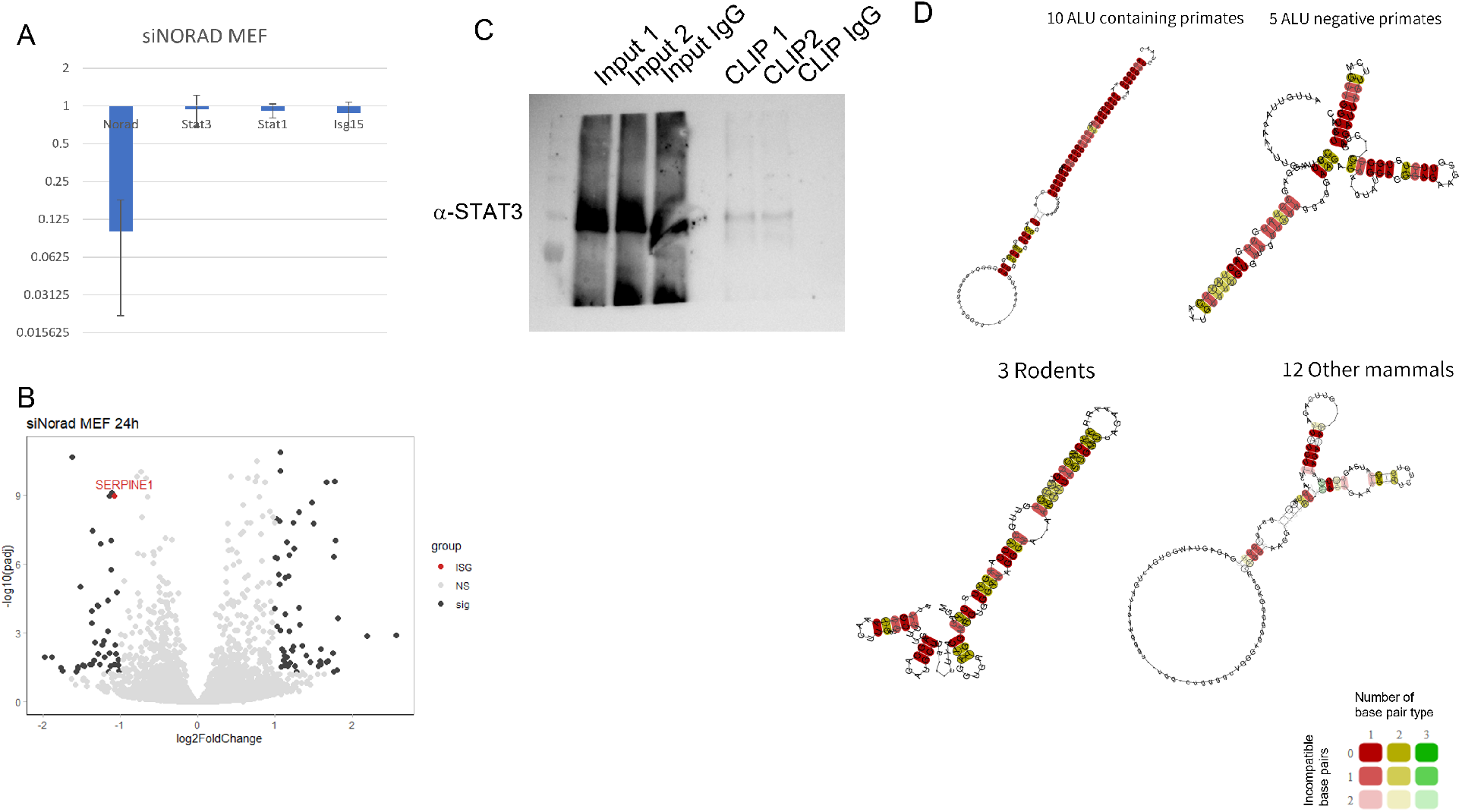
**(A)** qPCR analysis of Norad knockdown in MEF cells. Demonstrating that KD of Norad does not change the RNA levels of Stat3, Stat1, and Isg15 **(B)** A volcano plot of RNA-seq data from Norad KD versus control siRNA in MEF cells 24 hours post-transfection. Differentially expressed (Padj<0.05, -2>FC>2) ISGs are marked in red. **(C)** Western blot analysis of Stat3 CLIP-qPCR in MEF cells. 5% of the IP taken for the analysis **(D)** 2D representation of the sequences of the STAT3 binding sites on NORAD from different species. Sequences were folded using the LocARNA Alignment & Folding prediction tool. The structure with the highest conservation was predicted for sequences from ALU-containing primates, significantly higher than for sequences extracted from non-ALU primates, Rodents, and other Mammals.

## References

1. W. H. Hudson, E. A. Ortlund, The structure, function and evolution of proteins that bind DNA and RNA. Nat Rev Mol Cell Biol. 15, 749–60 (2014).

2. Y. Long, X. Wang, D. T. Youmans, T. R. Cech, How do lncRNAs regulate transcription? Sci Adv. 3 (2017), doi:10.1126/sciadv.aao2110.

3. S.-Y. Ng, R. Johnson, L. W. Stanton, Human long non-coding RNAs promote pluripotency and neuronal differentiation by association with chromatin modifiers and transcription factors. EMBO J. 31, 522–533 (2012).

4. Z. E. Holmes, D. J. Hamilton, T. Hwang, N. V. Parsonnet, J. L. Rinn, D. S. Wuttke, R. T. Batey, The Sox2 transcription factor binds RNA. Nat Commun. 11, 1805 (2020).

5. L. Hou, Y. Wei, Y. Lin, X. Wang, Y. Lai, M. Yin, Y. Chen, X. Guo, S. Wu, Y. Zhu, J. Yuan, M. Tariq, N. Li, H. Sun, H. Wang, X. Zhang, J. Chen, X. Bao, R. Jauch, Concurrent binding to DNA and RNA facilitates the pluripotency reprogramming activity of Sox2. Nucleic Acids Res. 48, 3869–3887 (2020).

6. S.-Y. Ng, G. K. Bogu, B. S. Soh, L. W. Stanton, The Long Noncoding RNA RMST Interacts with SOX2 to Regulate Neurogenesis. Mol Cell. 51, 349–359 (2013).

7. H. Luo, G. Zhu, M. A. Eshelman, T. K. Fung, Q. Lai, F. Wang, B. B. Zeisig, J. Lesperance, X. Ma, S. Chen, N. Cesari, C. Cogle, B. Chen, B. Xu, F.-C. Yang, C. W. E. So, Y. Qiu, M. Xu, S. Huang, HOTTIP-dependent R-loop formation regulates CTCF boundary activity and TAD integrity in leukemia. Mol Cell. 82, 833–851.e11 (2022).

8. H. Guo, J. Liu, Q. Ben, Y. Qu, M. Li, Y. Wang, W. Chen, J. Zhang, The aspirin-induced long noncoding RNA OLA1P2 blocks phosphorylated STAT3 homodimer formation. Genome Biol. 17, 24 (2016).

9. K. N. Smith, J. Starmer, T. Magnuson, Interactome determination of a Long Noncoding RNA implicated in Embryonic Stem Cell Self-Renewal. Sci Rep. 8, 17568 (2018).

10. P. Wang, Y. Xue, Y. Han, L. Lin, C. Wu, S. Xu, Z. Jiang, J. Xu, Q. Liu, X. Cao, The STAT3-Binding Long Noncoding RNA lnc-DC Controls Human Dendritic Cell Differentiation. Science (1979). 344, 310–313 (2014).

11. J. Zhang, Z. Li, L. Liu, Q. Wang, S. Li, D. Chen, Z. Hu, T. Yu, J. Ding, J. Li, M. Yao, S. Huang, Y. Zhao, X. He, Long noncoding RNA TSLNC8 is a tumor suppressor that inactivates the interleukin-6/STAT3 signaling pathway. Hepatology. 67, 171–187 (2018).

12. S. Dvir, A. Argoetti, C. Lesnik, M. Roytblat, K. Shriki, M. Amit, T. Hashimshony, Y. Mandel-Gutfreund, Uncovering the RNA-binding protein landscape in the pluripotency network of human embryonic stem cells. Cell Rep. 35, 109198 (2021).

13. S. Lee, F. Kopp, T.-C. Chang, A. Sataluri, B. Chen, S. Sivakumar, H. Yu, Y. Xie, J. T. Mendell, Noncoding RNA NORAD Regulates Genomic Stability by Sequestering PUMILIO Proteins. Cell. 164, 69–80 (2016).

14. A. Tichon, N. Gil, Y. Lubelsky, T. Havkin Solomon, D. Lemze, S. Itzkovitz, N. Stern-Ginossar, I. Ulitsky, A conserved abundant cytoplasmic long noncoding RNA modulates repression by Pumilio proteins in human cells. Nat Commun. 7, 12209 (2016).

15. F. Kopp, M. M. Elguindy, M. E. Yalvac, H. Zhang, B. Chen, F. A. Gillett, S. Lee, S. Sivakumar, H. Yu, Y. Xie, P. Mishra, Z. Sahenk, J. T. Mendell, PUMILIO hyperactivity drives premature aging of Norad-deficient mice. Elife. 8 (2019), doi:10.7554/eLife.42650.

16. M. M. Elguindy, J. T. Mendell, NORAD-induced Pumilio phase separation is required for genome stability. Nature. 595, 303–308 (2021).

17. M. Munschauer, C. T. Nguyen, K. Sirokman, C. R. Hartigan, L. Hogstrom, J. M. Engreitz, J. C. Ulirsch, C. P. Fulco, V. Subramanian, J. Chen, M. Schenone, M. Guttman, S. A. Carr, E. S. Lander, The NORAD lncRNA assembles a topoisomerase complex critical for genome stability. Nature. 561, 132–136 (2018).

18. J. Eggenberger, D. Blanco-Melo, M. Panis, K. J. Brennand, B. R. tenOever, Type I interferon response impairs differentiation potential of pluripotent stem cells. Proc Natl Acad Sci U S A. 116, 1384–1393 (2019).

19. X. Wu, V. L. Dao Thi, Y. Huang, E. Billerbeck, D. Saha, H.-H. Hoffmann, Y. Wang, L. A. V. Silva, S. Sarbanes, T. Sun, L. Andrus, Y. Yu, C. Quirk, M. Li, M. R. MacDonald, W. M. Schneider, X. An, B. R. Rosenberg, C. M. Rice, Intrinsic Immunity Shapes Viral Resistance of Stem Cells. Cell. 172, 423-438. e25 (2018).

20. A. B. Keenan, D. Torre, A. Lachmann, A. K. Leong, M. L. Wojciechowicz, V. Utti, K. M. Jagodnik, E. Kropiwnicki, Z. Wang, A. Ma’ayan, ChEA3: transcription factor enrichment analysis by orthogonal omics integration. Nucleic Acids Res. 47, W212–W224 (2019).

21. I. Sadzak, M. Schiff, I. Gattermeier, R. Glinitzer, l. Sauer, A. Saalmüller, E. Yang, B. Schaljo, P. Kovarik, Recruitment of Stat1 to chromatin is required for interferon-induced serine phosphorylation of Stat1 transactivation domain. Proceedings of the National Academy of Sciences. 105, 8944–8949 (2008).

22. V. Cimica, H.-C. Chen, J. K. Iyer, N. C. Reich, Dynamics of the STAT3 transcription factor: nuclear import dependent on Ran and importin-β1. PLoS One. 6, e20188 (2011).

23. M. Garcia-Moreno, M. Noerenberg, S. Ni, A. l. Järvelin, E. González-Almela, C. E. Lenz, M. Bach-Pages, V. Cox, R. Avolio, T. Davis, S. Hester, T. J. M. Sohier, B. Li, G. Heikel, G. Michlewski, M. A. Sanz, L. Carrasco, E. P. Ricci, V. Pelechano, I. Davis, B. Fischer, S. Mohammed, A. Castello, System-wide Profiling of RNA-Binding Proteins Uncovers Key Regulators of Virus Infection. Mol Cell. 74, 196-211.e11 (2019).

24. E. Wyler, K. Mösbauer, V. Franke, A. Diag, L. T. Gottula, R. Arsiè, F. Klironomos, D. Koppstein, K. Hönzke, S. Ayoub, C. Buccitelli, K. Hoffmann, A. Richter, I. Legnini, A. Ivanov, T. Mari, S. Del Giudice, J. Papies, S. Praktiknjo, T. F. Meyer, M. A. Müller, D. Niemeyer, A. Hocke, M. Selbach, A. Akalin, N. Rajewsky, C. Drosten, M. Landthaler, Transcriptomic profiling of SARS-CoV-2 infected human cell lines identifies HSP90 as target for COVID-19 therapy. iScience. 24, 102151 (2021).

25. E. Wyler, V. Franke, J. Menegatti, C. Kocks, A. Boltengagen, S. Praktiknjo, B. Walch-Rückheim, J. Bosse, N. Rajewsky, F. Grässer, A. Akalin, M. Landthaler, Single-cell RNA-sequencing of herpes simplex virus 1-infected cells connects NRF2 activation to an antiviral program. Nat Commun. 10, 4878 (2019).

26. K. M. Gao, A. G. Derr, Z. Guo, K. Nündel, A. Marshak-Rothstein, R. W. Finberg, J. P. Wang, Human nasal wash RNA-Seq reveals distinct cell-specific innate immune responses in influenza versus SARS-CoV-2. JCI Insight. 6 (2021), doi:10.1172/jci.insight.152288.

27. D. Butler, C. Mozsary, C. Meydan, J. Foox, J. Rosiene, A. Shaiber, D. Danko, E. Afshinnekoo, M. MacKay, F. J. Sedlazeck, N. A. Ivanov, M. Sierra, D. Pohle, M. Zietz, U. Gisladottir, V. Ramlall, E. T. Sholle, E. J. Schenck, C. D. Westover, C. Hassan, K. Ryon, B. Young, C. Bhattacharya, D. L. Ng, A. C. Granados, Y. A. Santos, V. Servellita, S. Federman, P. Ruggiero, A. Fungtammasan, C.-S. Chin, N. M. Pearson, B. W. Langhorst, N. A. Tanner, Y. Kim, J. W. Reeves, T. D. Hether, S. E. Warren, M. Bailey, J. Gawrys, D. Meleshko, D. Xu, M. Couto-Rodriguez, D. Nagy-Szakal, J. Barrows, H. Wells, N. B. O’Hara, J. A. Rosenfeld, Y. Chen, P. A. D. Steel, A. J. Shemesh, J. Xiang, J. Thierry-Mieg, D Thierry-Mieg, A. Iftner, D. Bezdan, E. Sanchez, T. R. Campion, J. Sipley, L. Cong, A. Craney, P. Velu, A. M. Melnick, S. Shapira, l. Hajirasouliha, A. Borczuk, T. Iftner, M. Salvatore, M. Loda, L. F. Westblade, M. Cushing, S. Wu, S. Levy, C. Chiu, R. E. Schwartz, N. Tatonetti, H. Rennert, M. Imielinski, C. E. Mason, Shotgun transcriptome, spatial omics, and isothermal profiling of SARS-CoV-2 infection reveals unique host responses, viral diversification, and drug interactions. Nat Commun. 12, 1660 (2021).

28. I. Ramos, G. Smith, F. Ruf-Zamojski, C. Martínez-Romero, M. Fribourg, E. A. Carbajal, B. M. Hartmann, V. D. Nair, N. Marjanovic, P. L. Monteagudo, V. A. DeJesus, T. Mutetwa, M. Zamojski, G. S. Tan, C. Jayaprakash, E. Zaslavsky, R. A. Albrecht, S. C. Sealfon, A. García-Sastre, A. Fernandez-Sesma, Innate Immune Response to Influenza Virus at Single-Cell Resolution in Human Epithelial Cells Revealed Paracrine Induction of Interferon Lambda 1. J V ol. 93 (2019), doi:10.1128/JVI.00559-19.

29. J. Ouyang, X. Zhu, Y. Chen, H. Wei, Q. Chen, X. Chi, B. Qi, L. Zhang, Y. Zhao, G. F. Gao, G. Wang, J.-L. Chen, NRAV, a Long Noncoding RNA, Modulates Antiviral Responses through Suppression of Interferon-Stimulated Gene Transcription. Cell Host Microbe. 16, 616–626 (2014).

30. H. Nishitsuji, S. Ujino, S. Yoshio, M. Sugiyama, M. Mizokami, T. Kanto, K. Shimotohno, Long noncoding RNA #32 contributes to antiviral responses by controlling interferon-stimulated gene expression. Proceedings of the National Academy of Sciences. 113, 10388–10393 (2016).

31. S. Shirahama, R. Onoguchi-Mizutani, K. Kawata, K. Taniue, A. Miki, A. Kato, Y. Kawaguchi, R. Tanaka, T. Kaburaki, H. Kawashima, Y. Urade, M. Aihara, N. Akimitsu, Long noncoding RNA U90926 is crucial for herpes simplex virus type 1 proliferation in murine retinal photoreceptor cells. Sci Rep. 10, 19406 (2020).

32. S. Agarwal, T. Vierbuchen, S. Ghosh, J. Chan, Z. Jiang, R. K. Kandasamy, E. Ricci, K. A. Fitzgerald, The long non-coding RNA LUCAT1 is a negative feedback regulator of interferon responses in humans. Nat Commun. 11, 6348 (2020).

33. O. Ziv, S. Farberov, J. Y. Lau, E. Miska, G. Kudla, l. Ulitsky, Structural features within the NORAD long noncoding RNA underlie efficient repression of Pumilio activity, doi:10.1101/2021.11.19.469243.

34. S. Will, T. Joshi, I. L. Hofacker, P. F. Stadler, R. Backofen, LocARNA-P: accurate boundary prediction and improved detection of structural RNAs. RNA. 18, 900–14 (2012).

35. J. M. Burke, E. T. Lester, D. Tauber, R. Parker, RNase L promotes the formation of unique ribonucleoprotein granules distinct from stress granules. J Biol Chem. 295, 1426–1438 (2020).

36. T. Matheny, B. Van Treeck, T. N. Huynh, R. Parker, RNA partitioning into stress granules is based on the summation of multiple interactions. RNA. 27, 174–189 (2021).

37. S. Namkoong, A. Ho, Y. M. Woo, H. Kwak, J. H. Lee, Systematic Characterization of Stress-Induced RNA Granulation. Mol Cell. 70, 175-187.e8 (2018).

38. I. Ulitsky, Evolution to the rescue: using comparative genomics to understand long noncoding RNAs. Nat Rev Genet. 17, 601–614 (2016).

39. Y. Guo, Utilization of different anti-viral mechanisms by mammalian embryonic stem cells and differentiated cells. Immunol Cell Biol. 95, 17–23 (2017).

40. T. Hashimshony, N. Senderovich, G. Avital, A. Klochendler, Y. de Leeuw, L. Anavy, D. Gennert, S. Li, K. J. Livak, O. Rozenblatt-Rosen, Y. Dor, A. Regev, I. Yanai, CEL-Seq2: sensitive highly-multiplexed single-cell RNA-Seq. Genome Biol. 17, 77 (2016).

41. M. Martin, Cutadapt removes adapter sequences from high-throughput sequencing reads. EMBnet J. 17, 10 (2011).

42. A. Dobin, C. A. Davis, F. Schlesinger, J. Drenkow, C. Zaleski, S. Jha, P. Batut, M. Chaisson, T. R. Gingeras, STAR: ultrafast universal RNA-seq aligner. Bioinformatics. 29, 15–21 (2013).

43. S. Anders, P. T. Pyl, W. Huber, HTSeq—a Python framework to work with high-throughput sequencing data. Bioinformatics. 31, 166–169 (2015).

44. M. I. Love, W. Huber, S. Anders, Moderated estimation of fold change and dispersion for RNA-seq data with DESeq2. Genome Biol. 15, 550 (2014).

45. W. Walter, F. Sánchez-Cabo, M. Ricote, GOplot: an R package for visually combining expression data with functional analysis. Bioinformatics. 31, 2912–2914 (2015).

46. M. D. Luecken, F. J. Theis, Current best practices in single-cell RNA-seq analysis: a tutorial. Mol Syst Biol. 15 (2019), doi:10.15252/msb.20188746.

47. T. Ilicic, J. K. Kim, A. A. Kolodziejczyk, F. O. Bagger, D. J. McCarthy, J. C. Marioni, S. A. Teichmann, Classification of low quality cells from single-cell RNA-seq data. Genome Biol. 17, 29 (2016).

48. A. T. L. Lun, D. J. McCarthy, J. C. Marioni, A step-by-step workflow for low-level analysis of single-cell RNA-seq data with Bioconductor. F1000Res. 5, 2122 (2016).

49. O. Gascuel, BIONJ: an improved version of the NJ algorithm based on a simple model of sequence data. Mol Biol Evol. 14, 685–695 (1997).

50. I. Letunic, P. Bork, Interactive tree of life (iTOL) v3: an online tool for the display and annotation of phylogenetic and other trees. Nucleic Acids Res. 44, W242–W245 (2016).

